# Elucidating changes in gene expression driving nuclear dynamics across a foraminiferal life cycle with Pheno2Seq

**DOI:** 10.64898/2026.10.21.760831

**Authors:** Elinor G. Sterner, Giulia M. Ribeiro, Marie Leleu, Laura A. Katz

## Abstract

In many microeukaryotes, conserved morphology masks transitions between haploid and diploid phases of the life cycle, making it difficult to link gene expression to development. Hence, insights on such transitions come largely from ‘model’ lineages of animals, plants and fungi. To address this gap, we developed a method, Pheno2Seq, that enables visualization of fluorescently-stained individuals followed by identification of candidate genes through single-cell transcriptome analyses. We applied this method in *Allogromia laticollaris*, a foraminifera with a complex life cycle that alternates between haploid and diploid phases and includes endoreplication to ∼11,000N. As this taxon lacks a reference genome, we used a conservative phylogenomic approach to identify gene families present across diverse foraminifera. We identified >800 differentially expressed genes underlying life cycle transitions along with a similar number of constitutively-expressed genes. During *Zerfall*, a phase of extreme endoreplication, the transcriptome is largely silenced with the exception of gene families involved in genome processes and stability. Intriguingly, these gene families include chromatin modifiers, spliceosomal components and histone paralogs, some of which are known to contribute to cell type differentiation in multicellular eukaryotes. We also find numerous genes involved in DNA repair (e.g. DNA damage response and R-loop regulating genes). We speculate that these genes are part of an ancient eukaryotic toolkit, likely present in the last eukaryotic common ancestor (LECA) and that regulates genomes during life cycle transitions. Our method, Pheno2Seq, can be adapted to diverse lineages to enable the study of life cycle regulation in microeukaryotes that lack morphologically distinguishable life cycle stages.

**Significance:** Unicellular eukaryotes represent most of the diversity across the eukaryotic tree of life, but life cycle transitions in many of these species can be cryptic (i.e. multiple stages share identical morphology despite changing nuclear phenotypes). We develop Pheno2Seq (Phenotype to Sequence), which characterizes nuclear phenotype by fluorescence microscopy, then elucidates gene expression underlying nuclear stage transitions. Changes in nuclear phenotype in the foraminifera sampled here demonstrates a central role of the reprogramming of histones and other genes involved in chromatin regulation, suggesting that the diverse life cycles observed across the eukaryotic tree of life emerged from a shared toolkit present in the last eukaryotic common ancestor (LECA).

## Introduction

Changes in nuclear architecture underlie life cycle transitions, yet our understanding of such changes is largely limited to a few model species of animals and plants. Analyses of microbial eukaryotes (i.e. protists) have uncovered life cycles marked by dynamic genome processes that do not fit into standard models. This is exemplified by extensive chromosomal rearrangements in ciliates, programmed karyotype instability in some parasitic protists and dramatic shifts in ploidy and genome organization in foraminifera (reviewed in 1). Protist life cycle transitions are often cryptic, with multiple life history stages indistinguishable by light microscopy (e.g., 2–4).

Even in the absence of morphological changes, alterations in nuclear phenotype indicate significant genome modifications including separation of germline and somatic DNA in ciliates and some foraminifera (5, 6), sexual and parasexual reproduction (7), cyclic polyploidy with chromatin extrusion in amoebae (8) and extreme endoreplication that couples nuclear decay in some foraminifera (9).

Even in lineages with well-described life cycles, changes in gene expression driving transitions are largely unknown, with most data coming from better-studied parasites plus lineages for which cultures can be synchronized. Across diverse eukaryotes, histones are modified or exchanged for variant histones to regulate life cycle transitions, as has been demonstrated in *Arabidopsis* (10), humans (11), *Trypanosoma cruzi* (12)*, Toxoplasma gondii* (13), *Plasmodium falciparum,* the causative agent of malaria (14), and many other species (15). In ciliates, a clade in which distinct germline and somatic nuclei coexist in every single-celled individual, molecular mechanisms for restructuring nuclei and genomes include regulation of both chromatin modifiers and small RNAs pathways (e.g. dicer-like and argonaute proteins, 16). Though a limited amount of data on differential gene expression exists on less-well studied lineages like brown and red algae (17, 18), most protist lineages remain understudied, in part because of challenges working with species with cryptic life cycle stages.

We investigate the foraminiferal species *Allogromia laticolaris* to probe gene expression changes that underlie life cycle transitions in a species with a cryptic alternation between haploid and diploid generations. Foraminifera are a group of predominantly marine shell-building amoebae that are critical components of marine food webs (19). Those with calcareous tests are biomarkers for past climatic conditions in the fossil record (e.g. 20). Due to their relatively large size and abundance, foraminifera have been the subject of morphological study nearly since the invention of the microscope (reviewed in 21), revealing that the complex life cycles of foraminifera are marked by extreme changes in genome organization (5). A typified foraminifera life cycle is marked by alternation of haploid and diploid generations, which is rarely observed outside of plants. Certain species of foraminifera have documented distinct germline and somatic nuclei (5), another indication of the genome dynamics in this lineage.

The life cycle of *A. laticollaris* has been studied using light and fluorescence microscopy (9, 22, 23). Like land plants, foraminifera have a complex alternation of generations, though with some notable twists: multinucleated diploid stages go through meiosis to produce uninucleate stages that ultimately produce haploid gametes, which in turn fuse to generate multinucleated diploid phases (5, 24). Foraminifera like *A. laticollaris* reach large sizes by endoreplicating their genomes, with an estimated 11,000 gene copies present in large uninucleate individuals (9).

These large individuals eventually undergo nuclear decay in a process termed *Zerfall*, which generates hundreds of gametic nuclei that next fuse within the parent (9, 24, 25). Cell fission generates multinucleated offspring within the parental test, or schizont, before juveniles emerge to develop into multinucleated diploid phases, thus restarting the life cycle (9, 22, 23).

Here we evaluate gene expression changes underlying cryptic life cycle transitions in *A. laticollaris.* As haploid uninucleate and diploid multinucleate life cycle stages look nearly identical by light microscopy, it is very challenging to link gene expression patterns with developmental stages (Fig. 1A). Thus, we developed Pheno2Seq, a method that combines fluorescence microscopy with single cell transcriptomics, allowing analysis of differential gene expression across morphologically cryptic life cycle stages. We then analyze the resulting data using Eukphylo (26), a taxon-rich phylogenomic pipeline, to identify gene families and separate sequences belonging to *A. laticollaris* from contaminants. Finally, we explore the resulting data using GO terms (27) as well as by mining the literature for candidate gene families involved in genome reorganization. The resulting data provide a first look at differential gene expression in foraminifera while also offering a novel method, Pheno2Seq, that will be of use to researchers studying the many other protist lineages that are marked by cryptic life cycle stages.

**Fig. 1.**
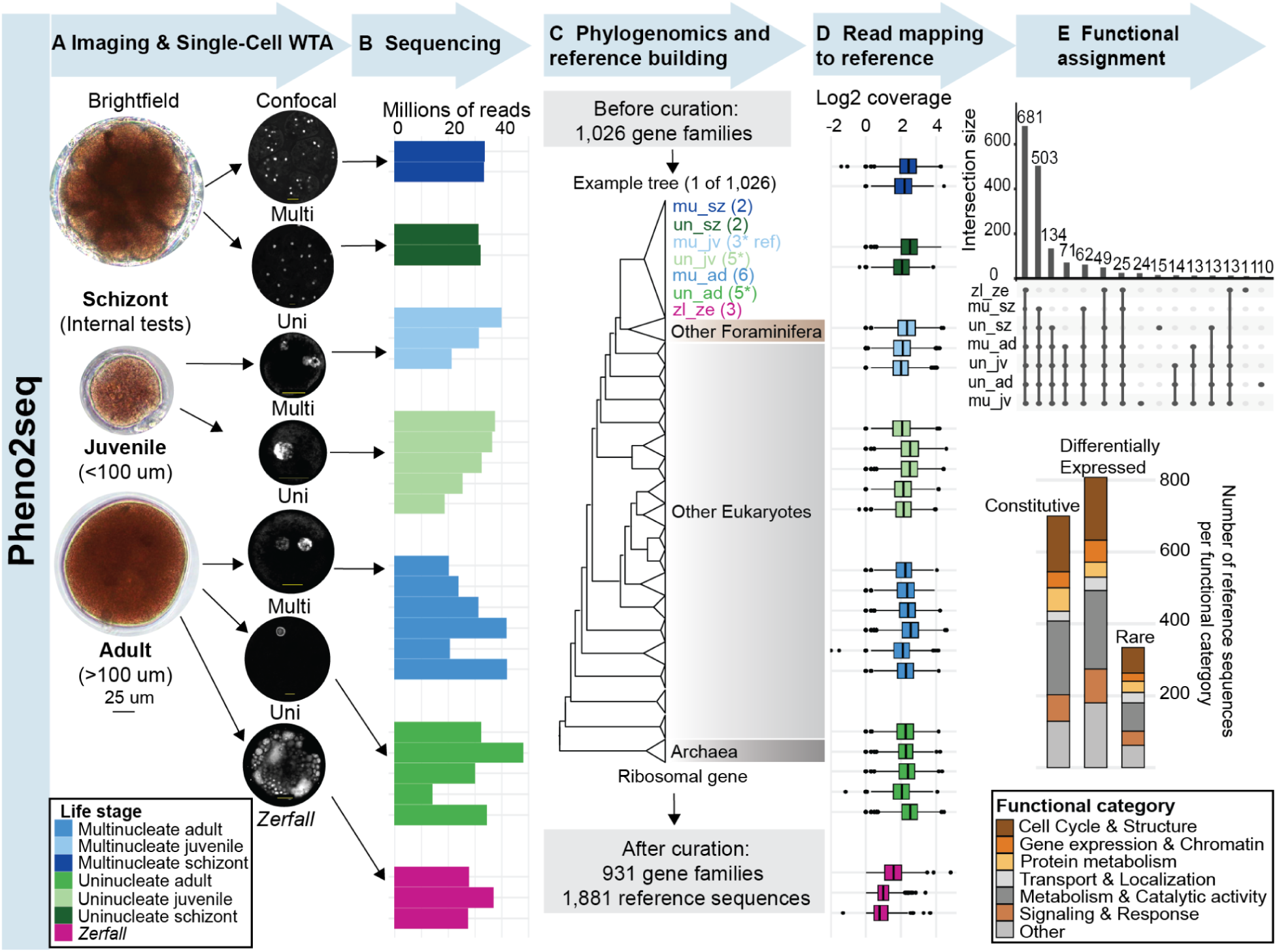
Workflow of Pheno2Seq enables characterization of nuclear phenotype followed by associated gene expression changes. **(A)** Light microscopy only allows characterization of *A. laticollaris* individuals as schizonts (i.e. cells undergoing fission), juveniles, and adults. Confocal microscopy following fixation with glyoxal enables characterization of nuclear phenotypes (i.e. uninucleate, multinucleate, in *Zerfall*). **(B)** Following fixation, staining, and imaging, we sequenced transcriptomes from 26 cells sampled across seven life cycle stages. **(C)** Following gene family assignment and tree building with EukPhylo (26), we select *A. laticollaris* sequences to create a reference of 1,881 sequences from 1,026 gene families. **(D)** Even though cells had been (briefly) exposed to a UV laser, we found a similar number of reads per cell mapping to the reference. A notable exception were *Zerfall* cells, which we found have a very distinctly expressed transcriptome. **(E)** *top* Functional assignment reveals most gene families (681 sequences) are shared across all stages, or between all stages except *Zerfall* (503 sequences). **(E)** *bottom* Following differential gene expression analysis, we find similar distributions of functional categories in 708 constitutive genes, 825 differentially expressed genes, and 348 rare genes.

## Results and Discussion

### Pheno2seq enables single-cell transcriptome analysis following characterization of genome architecture

We developed Pheno2Seq, a method that integrates characterization of nuclear architecture by fluorescence confocal microscopy followed by single-cell transcriptomics and phylogenomic analysis of gene expression patterns across life cycle stages (Fig. 1). We analyzed 26 *A. laticollaris* individuals after determining the life history stage with laser scanning confocal microscopy (LSCM) of Hoechst-stained cells (Fig 1, Table S1). A novel aspect of Pheno2Seq is the fixation of cells in 1% glyoxal, which enables both staining of nuclei with Hoechst and subsequent characterization of transcriptome sequences. Glyoxal crosslinks proteins while leaving nucleic acids unattached, allowing for downstream amplification of transcripts (28, 29, e.g. 30). To our knowledge, this fixative has only been used in animal tissues, where it results in transcript quality comparable to unfixed tissues, albeit without exposure to UV light through fluorescence microscopy (30).

Fluorescence microscopy identified the life cycle stage (i.e. uninucleate vs multinucleate) of 26 individuals (Fig. 1A). This analysis yielded four schizonts (daughter cells contained in a parental test), eight juveniles, and 14 adults (Fig. 1A and B, dataS1A). Among the adults, we captured sufficient RNA from three cells in *Zerfall*, a process by which cells transition from uninucleate to multinucleate phases through a radical reorganization that includes a nucleus-free stage where DNA is distributed throughout the cytoplasm (Fig. 1A and B) (9). Interestingly, cDNA yields following single-cell whole transcriptome amplification varied by life stage, with the lowest yield being among the three *Zerfall* cells that yielded sufficient RNA (transcriptome amplification of six other *Zerfall* cells failed), which suggests transcription may be shut down during the major remodeling of nuclear architecture in *Zerfall*. Despite this variance, we obtained an average of 30 million reads per cell, enabling robust analysis of differential gene expression (Fig. 1B, data S1A).

To characterize gene expression, we used EukPhylo (26) to build 1,026 gene trees that include our 26 cells plus sequences from 944 outgroups (i.e. other foraminifera plus diverse eukaryotes and prokaryotes, Fig.1C, data S1D). These trees topologically distinguish host from contaminating sequences. Because *A. laticollaris* lacks a reference genome, we conservatively curated *A. laticollaris* clades from the ‘parental trees’ (data S2) to only retain those with sequences sister to other foraminifera, resulting in 41,137 sequences. Clustering the resulting sequences generated 1,881 reference transcriptome sequences, only four of which failed to meet minimum coverage requirements and were dropped from subsequent analyses (see methods, Fig. 1C, data S3).

Mapping reads from transcriptomes of 26 individuals to the reference yielded consistently high coverage across most life cycle stages (Fig. 1D). Despite the reduced transcription in *Zerfall,* we found a large number of shared gene families across all stages (681 reference sequences) with the second largest intersection of shared genes being present in all stages except *Zerfall* (503 reference sequences; Fig. 1E, data S4). Dividing genes into three categories identified 708 genes (38% of references) constitutively expressed genes (present in at least 80% of cells, and with no significant changes across life history stages) plus 348 (18.5% of references) ‘rare’ or lowly expressed genes (Fig. 1E). As discussed in the next section, 825 genes (46% of the reference) are differentially expressed, defined as having significant log fold changes in DESeq2 (see methods). Functional assignments using GO term annotations (27) are broadly similar across all three gene categories, though differentially expressed genes show proportionally greater contributions of cell cycle and chromatin related functions (Fig. 1E). In sum, Pheno2Seq addresses a knowledge gap in foraminifera (and many microeukaryotes) by providing the first analyses of changes in gene expression throughout their complex and cryptic life cycles.

Though research evaluating differential gene expression (DGE) in non-model organisms is increasing, DGE still faces multiple challenges including unculturability or asynchronous cultures, small cell sizes, and cryptic diversity (reviewed in 31). One strategy that has been employed is to cluster the cells by transcriptome profile similarity during analysis (32), but this loses a link with morphology. Pheno2seq offers a valuable new method to link fluorescence images of single cells to analyses of transcriptomes; moreover, Pheno2Seq could be used to image individuals with distinct morphologies from species that lack synchronous cultures, or for individuals isolated from different environments or seasons.

### Gene expression differs most between stages of major nuclear reorganization

After visualization of nuclear architecture by Hoechst-staining of cryptic life cycle stages, we analysed differential gene expression. After identifying gene families that belong to *A. laticollaris* with EukPhylo (26), we used DESeq2 (33, 34) to compare consecutive transitions (e.g. from uninucleate adults to *Zerfall,* and then from *Zerfall* to multinucleate schizonts; Fig. 2, data S4, see methods) and visualization on volcano plots illustrates the magnitude of changes at each transition (Fig. 2). Gene expression differs most dramatically as cells enter and exit *Zerfall* (Fig. 2). There are 492 differentially expressed genes at the transition from uninucleate adult to *Zerfall* (274 up-regulated in *Zerfall* and 218 up-regulated genes in uninucleate adults) and 539 differentially expressed genes at the transition between *Zerfall* and the multinucleate schizont (288 up-regulated in *Zerfall* and 251 genes up-regulated in multinucleate schizonts; Fig. 2, dataS1C). A dramatic reprogramming of gene expression is consistent with cytological observations of *Zerfall* during which the nuclear envelope breaks down and chromatin disperses throughout the cytoplasm, followed by the formation of gametes, then zygotes (through autogamy), and ultimately schizogony (i.e. cell fission) to generate multinucleate juveniles (9).

**Fig. 2.**
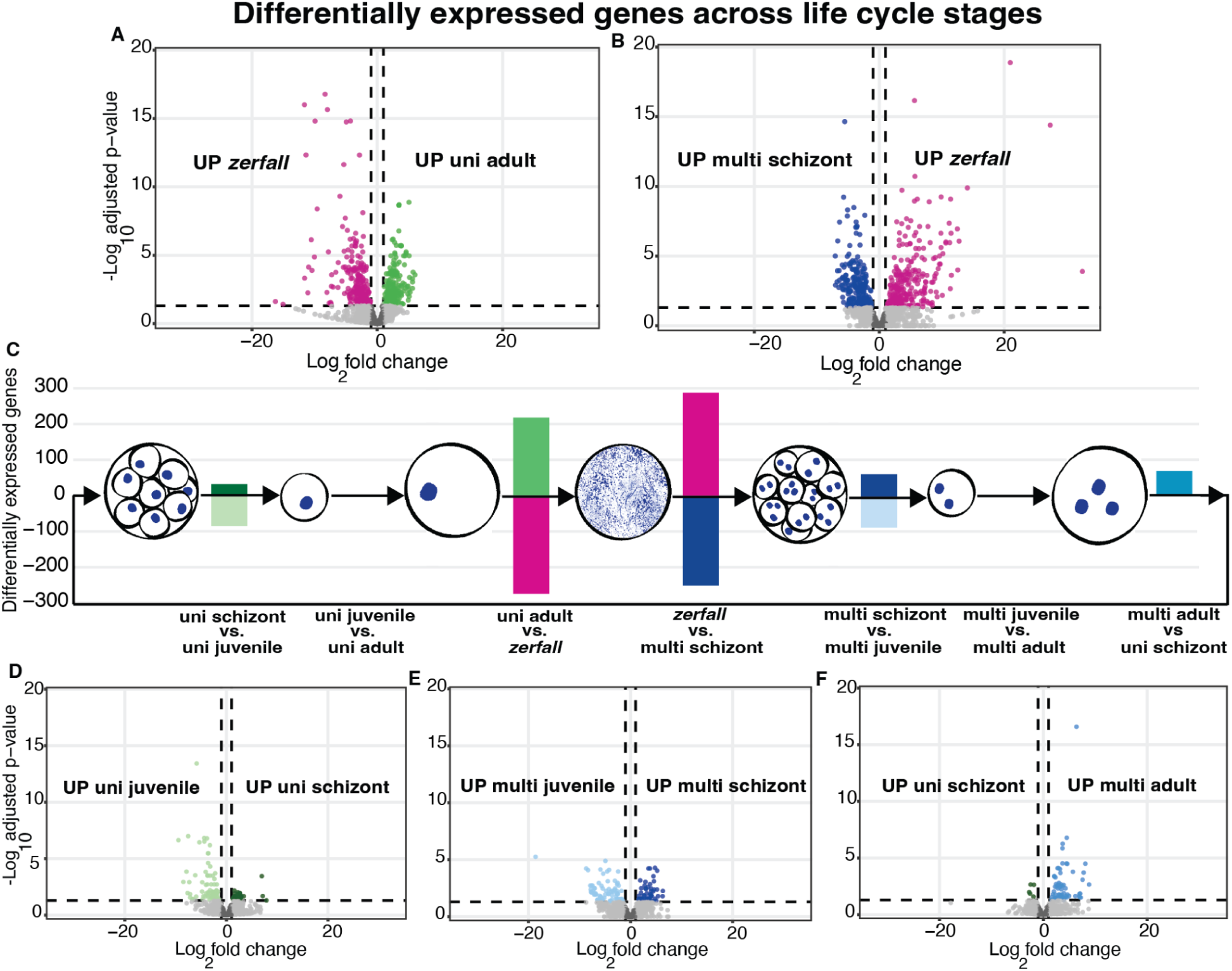
Pheno2Seq demonstrates that the greatest number of differentially expressed genes in the *Allogromia laticollaris* life cycle occur as the cell enters and exits *Zerfall*. Comparisons across consecutive life cycle stages show that genes are significantly altered as cells **(A)** enter and then **(B)** exit *Zerfall* during **(C)** the life cycle of *A. laticollaris.* In contrast, transitions that do not involve *Zerfall* have very few differentially expressed genes **(D-F),** particularly during growth phases. All comparisons are pairwise between two adjacent life cycles based on description in Timmons et al. (9). Life cycle colors are the same as in Fig. 1. Illustrations by Tejasvi Kumaran (Smith College)

As one example, the most upregulated gene in *Zerfall* is an arginine kinase (EC 2.7.3.2, data S8A), which has a normalized base mean of 175, 561 reads in *Zerfall* (compared to 121 in adults), representing a log2 fold change of 11.62. We propose this protein is used as an efficient energy reserve during *Zerfall*, a period of reduced transcription; the role of arginine kinases as an energy source is well documented (35), and seen in other protists including *Phytophthora,* the causative agent of the Irish potato famine (36), and *Tetrahymena* (37).

In contrast to the dramatic shifts in gene expression as cells enter and exit *Zerfall*, we found only one differentially expressed gene when multinucleate and uninucleate juveniles transition to adults (Fig 2). These data are consistent with other organisms where continuous developmental trajectories (i.e. growth) do not correspond to significant changes in gene expression. Like *A. laticollaris,* similar patterns of limited transcriptional contrast between growth transitions have been reported in the development of butterflies, nematodes, and water fleas (38). In contrast to growth stages, moderate reprogramming happens when *A. laticollaris* cells enter reproductive schizont stages. In our comparisons, we found 149 differentially expressed genes between multinucleate schizonts and multinucleate juvenile, 71 between multinucleate adult and uninucleate schizont, and 117 between uninucleate schizonts and uninucleate juveniles (Fig. 2, data S1C), with the majority of these genes related to translation (Fig. 3). The moderate reprogramming of gene expression that we observe within schizont stages are consistent with analogous changes in some animals such as in the crustacean *Daphnia magna* (38).

**Fig. 3.**
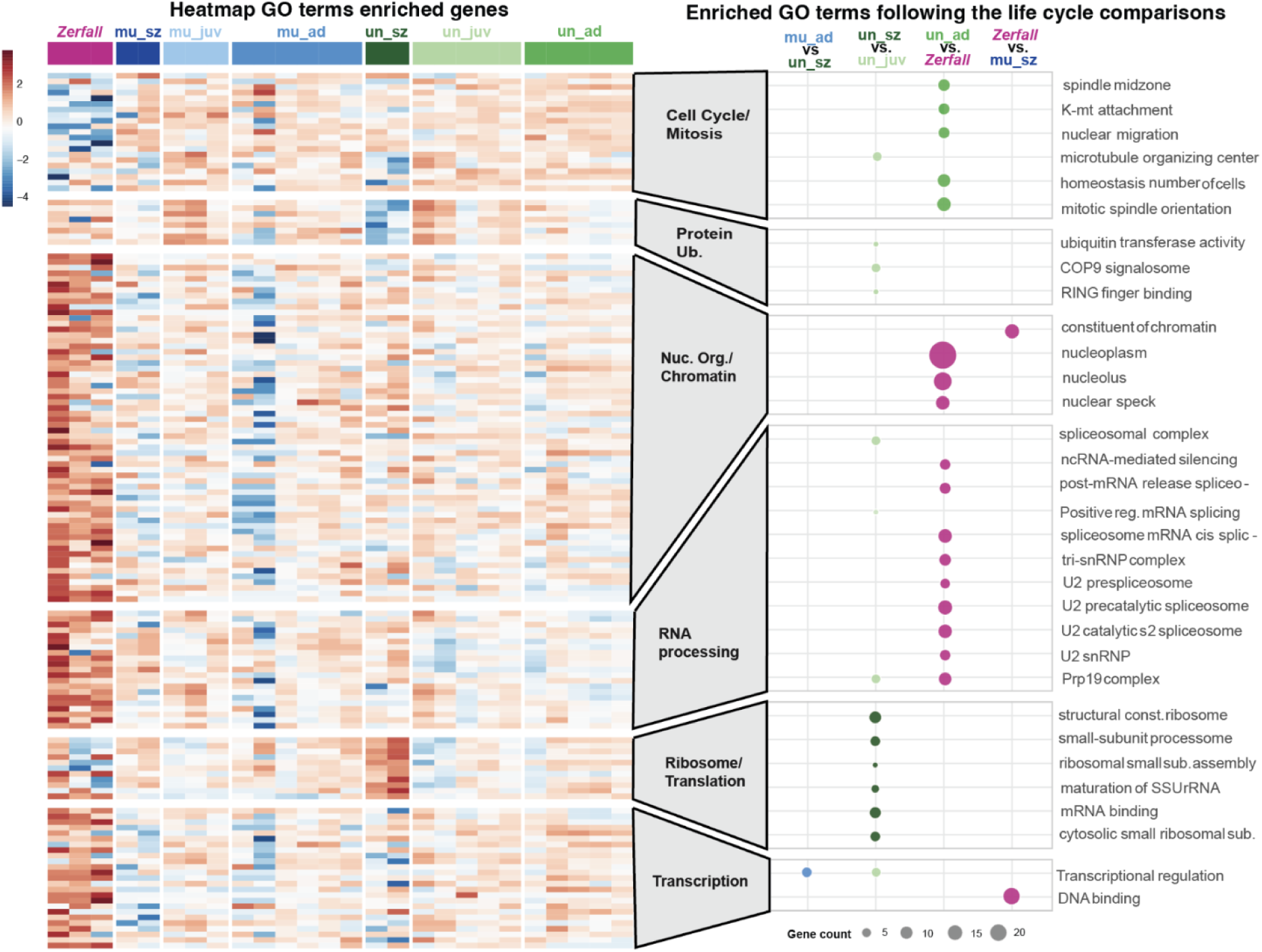
GO term enrichment analysis of life cycle transitions demonstrates the role of chromatin modifiers and spliceosomal components underlying the dramatic restructuring during *Zerfall*. Left: Heatmap of differentially expressed genes across life cycle stages (upper legend) clustered by functional (GO) category. Right: Enriched GO terms associated with genetic processes include terms associated with cell cycling, protein ubiquitination, nuclear reorganization, RNA processing (i.e. the spliceosome), and both transcription and translation. Life cycle colors are the same as in Fig. 1.

While *Zerfall* is unique to foraminifera, similar nuclear changes in nuclear architecture are found in other eukaryotes. Across diverse protists, chromatin condensation, nuclear envelope remodeling, and programmed DNA elimination delineate some of the transcriptional boundaries in life cycles (16, 39). In the case of *Candida albicans,* polyploidy and other forms of genetic instability generate genetic diversity, and can be driven by aberrant histone marks (e.g., H3K56ac) or failed cytokinesis during external stress (40). In some animals, endoreplication occurs in healthy cells (e.g. *Drosophila* larvae) and requires suppression of normal cell cycle regulatory genes, particularly Cdk1 (41). In *Allogromia*, the story is more complicated as one of three copies of Cdk1 (OG6_100334) is upregulated in *Zerfall*, while two others are higher in multinucleated schizonts (DataS8A). This capacity for cells to grow and generate genetic diversity at risk of genetic instability is found across the eukaryotic tree of life (1). In sum, Pheno2Seq demonstrates a similar pattern in *A. laticollaris* as compared to other eukaryotes that remain transcriptionally stable during the vegetative growth followed by pronounced transcriptional remodeling during transitions that involve changes to their nuclear architecture (16, 42).

### Functional analyses of differentially expressed genes highlight roles of cell cycle components, chromatin modifiers and the spliceosome

We explored function through analysis of gene ontology (GO Terms, 27) to assign to the differentially expressed genes between life cycle stages, categorizing genes into six distinct clusters across the seven life cycle stages (Fig. 3, data S1E). Enrichment analysis (Fig. 3, right panel), yielded terms related to cell cycle and mitosis (e.g. spindle midzone, kinetochore–microtubule attachment, nuclear migration) as most strongly enriched in the conditions down regulated as cells enter *Zerfall* (16 differentially expressed transcripts, several of them corresponding to Cdc42; DataS8A), which suggests that *Zerfall* is suppressing the cellular machinery required for nuclear division. This is consistent with the nuclear breakdown (9) and reduction in transcription (this study) associated with *Zerfall*. As uninucleate schizonts transition into uninucleate juveniles, genes associated with protein ubiquitination and quality control are upregulated (e.g. five heat shock proteins, one ubiquitin ligase), leading to an enrichment of GO terms including ubiquitin transferase activity, COP9 signalosome and RING finger binding (Fig. 3, DataS8A). Some of these transcripts are also upregulated other life cycle transitions including *Zerfall* (e.g. ubiquitin ligase, RING finger binding protein, and one heat shock protein) and the transition to multinucleated juveniles (e.g. ubiquitin protein ligase). Similar to *A. laticollaris,* ubiquitin proteins are involved in life cycle transitions in other eukaryotes including the slime mold *Dictyostelium discoideum* (43), the pathogens *Acanthamoeba* and *Entamoeba* (44), as well as in plants (45), animals (46) and fungi (47).

In addition to transcriptional changes in cellular processes, we find evidence for upregulation of genes related to chromatin and the spliceosome during *Zerfall*, a life cycle transition where ∼11k genome copies are processed to generate gametic nuclei (9). Gene families enriched in *Zerfall* include nuclear-associated GO terms (e.g., chromatin, nucleoplasm, nucleolus, DNA binding and RNA processing terms), as well as spliceosomal components (e.g., U2 related terms, Prp19 complex; Fig 3, data S1E). The upregulation of spliceosomal terms is consistent with either: 1) post-transcriptional regulation of genes expressed during *Zerfall*, or 2) non-splicing functions of the spliceosome. For the latter, there is a growing literature on the plurality of functions of the spliceosome, where components are involved in chromatin modification and double-strand break repair (48–50), processes that may be ongoing during *Zerfall*. For example, the splicing factor PRP19 has been associated with DNA repair by detecting its presence at single strand DNA breaks (48) while proteins like BUD31 and SNRPA are recruited to regions of double-strand breaks (50). As cells exist *Zerfall* and juveniles emerge from uninucleate schizonts, they upregulate genes enriched in ribosomal and translation-related GO terms (Fig. 3, data S1E), consistent with increased translational demand as cells prepare for division and growth.

Next, we identified several candidate genes relevant to DNA repair and genome stability. Intriguingly, we found numerous genes upregulated in *Zerfall* relevant to R-loop regulation. R-loops are non-B DNA structures involving single-stranded RNA and double-stranded DNA; though they typically result from transcription, they also can form during DNA replication with the introduction of Okazaki fragments (51). We propose that R-loop associated genes may play a role in the dramatic rearrangement of the genome during *Zerfall*. Notably, *A. laticollaris* upregulates several gene families associated with R-loops including DHX family RNA helicases, topoisomerase (Top1), the methyltransferase METTL3 (51) and multiple sirtuins. We found numerous upregulated gene families associated with DNA repair and genome stability (data S8A) including numerous DNA helicases (MCM family (52)), DNAJ (50), SMC family genes (51, 52), as well as the DNA replication enzyme GINS (52). Finally, *Zerfall* cells upregulate PIWI, a protein involved in both chromatin stability and germline/soma distinctions in diverse eukaryotes (53–55). Though there are many steps ahead to demonstrate robustly the roles of these proteins during *Zerfall* (i.e. transforming *A. laticollaris* into a tractable genetic system for gene knock out), Pheno2Seq has generated a robust list of candidate genes.

Genes identified by Pheno2Seq as underlying the dramatic developmental changes in *A. laticollaris* (Fig. 3; data S1E) are consistent with observations from distantly related eukaryotic clades including plants and animals, and provide evidence for a conserved eukaryotic toolkit for genome remodeling (56, 57). In animals, changes in nuclear architecture underlie life cycle transitions (58), for example during embryonic development of both mammals (59) and zebrafish (60). Plants also rely on functionally similar genes during gametogenesis (61), and chromatin reprogramming mechanisms include switching between histone variants (62). Hence, the chromatin and nuclear architecture changes we observe as *A. laticollaris* appear to rely on a toolkit shared with diverse eukaryotes, illuminating mechanisms at LECA.

### Life cycle transitions are also marked by differentially expressed paralogs of histones

We identified differences in expression patterns within histone families, and particularly among paralogs of histone H4 and H2A. Notably, such changes are not evident in differential gene analyses as they represent changes within a gene family (i.e. among paralogs) rather than between gene families. To investigate differential expression of paralogs, we first inspected readmapping data for references assigned to the core histones (H2A, H2B, H3, H4; data S1C; Fig. 4, top left). To our surprise, we found that *A. laticollaris* appears to have 17, 8, 7, and 7 divergent paralogs of histones H2A, H2B, H3 and H4, respectively (Fig. 4, SFigure1, data S2). Roughly half of the histone paralogs are differentially regulated in at least one life stage comparison, and several are *Zerfall* specific (H2A 9/18 and H4 4/8; Figure 4). Our phylogenetic analysis of histone H4 identified six ancient clades, several of which show differential expression patterns (Fig. 4; data S2). H4 transcripts in clade 2 are expressed in all stages, but have a lower abundance in *Zerfall* cells while H4 transcripts in clades 3-6 are only found in uninucleate and *Zerfall* cells. This pattern is consistent with a gradual increase of the expression of many H4 paralogs as the cell enters *Zerfall*, followed by a drop in expression once multinucleate schizonts are formed (Fig 4F). Histone H2A also has divergent paralogs; we find seventeen paralogs falling in distinct foraminiferal clades. In two of these H2A paralogs (clades 1a and 1b, SFigure1, data S1C) the sequences are substantially upregulated in *Zerfall* while other paralogs are upregulated in transitions in and out of the uninucleate schizont stage (Fig 4).

**Fig. 4.**
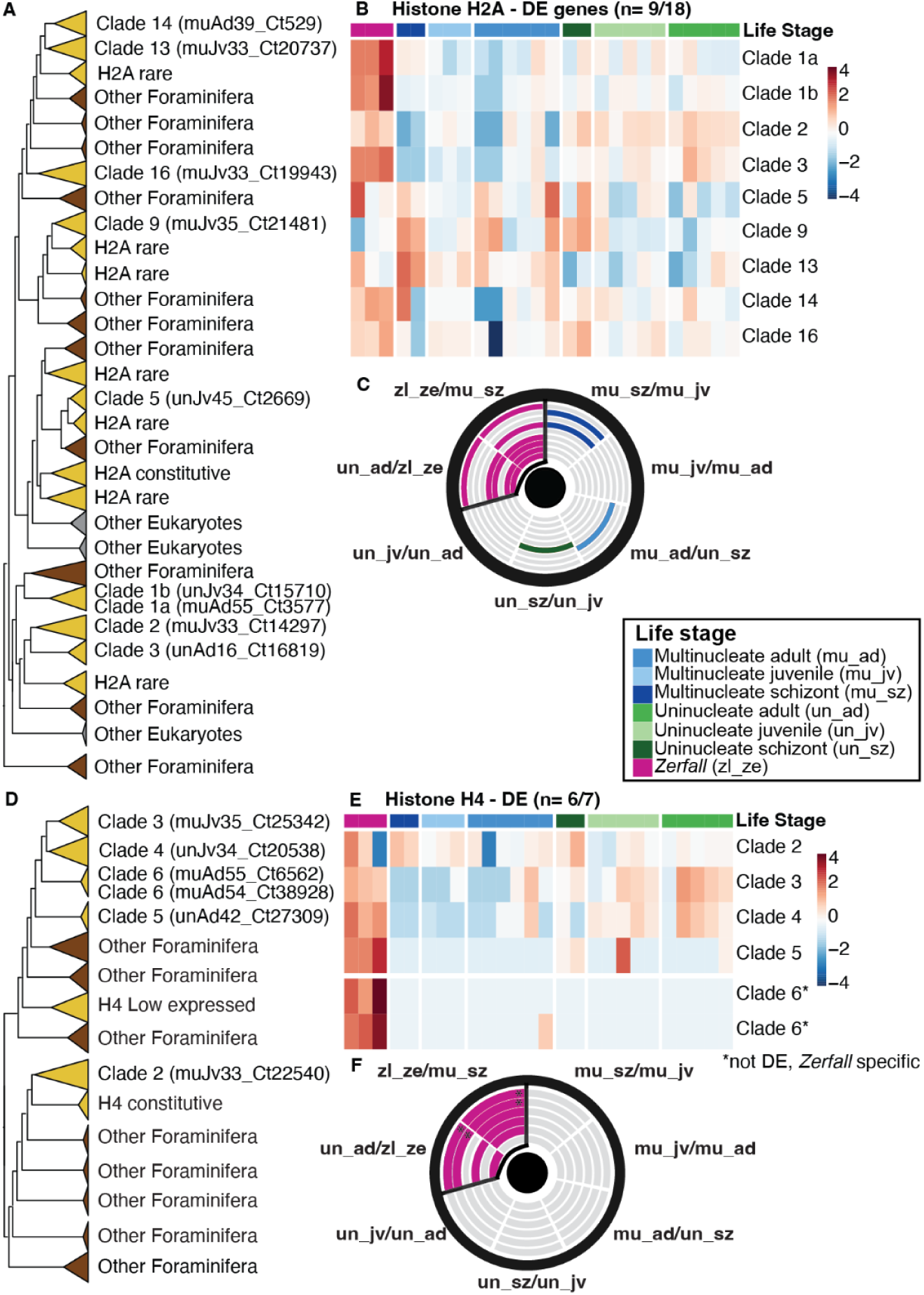
Histones H2A and H4 paralogs show dramatic shifts as cells enter and exit *Zerfall*. **(A)** Histone H2A (OG6_100077) gene tree with divergent paralogs in *A. laticollaris marked* in yellow. **(B)** Heatmap of all differentially expressed Histone H2A paralogs, and **(C)** Coulson plot showing differential expression of H2A paralogs that are mostly upregulated in *Zerfall* (pink stripes within circle), with a few paralogs upregulated in other stages. **(D)** Histone H4 (OG6_100120) gene tree, and **(E)** heatmap of all differentially expressed Histone H4 paralogs and two *Zerfall-*specific clades. **(F)** Coulson plot showing differential expression of H4 paralogs are mostly upregulated in *Zerfall* (pink stripes within circle). In both Coulson plots, the paralogs are arrayed from inside to out following the top-to-bottom arrangement on the heatmap. The “*” indicates the *Zerfall-*specific clades that were not statistically differentially expressed in the pairwise comparisons.

Differential expression of histone paralogs are associated with life cycle transitions and chromatin modifications in diverse eukaryotes. For example, transitions between asexual and sexual life stages in *Plasmodium falciparum* rely on the differential regulation of histone variants to structure chromatin between life stages (63). Life cycle transitions in a number of taxa have also been associated with differential regulation of histone H2A variants. For example, histone H2A variants are associated with transcriptional initiation and DNA repair in animals (64), regulation of TEs in plants (65), transcriptional regulation in response to host shifts in *Toxoplasma* (66) and chromatin modifications in ciliates (67). While core histones are generally expressed and incorporated into chromatin only during the S phase, variants are exchanged by chaperone proteins throughout the life cycle (67, 68), consistent with our findings in *A. laticollaris.* The ubiquity of life cycle-specific histone variants indicate that LECA may have expanded its histone gene families, facilitating the evolution of diverse life cycles.

### Synthesis

Pheno2Seq effectively combines identification of nuclear phenotype with analyses of gene expression, essential for studying species like *A. laticollaris* for which life cycle stages are cryptic. Throughout alternation of generations and extreme changes in ploidy (1 - 11,000N), *A. laticollaris* differentially regulates expression of spliceosomal components, genes related to chromatin management and histone variants (especially H2A). Together these data suggest that *A. laticollaris* relies on a toolkit of genes shared across diverse eukaryotes (e.g. among plants, animals, fungi, ciliates, trypanosomes, *Plasmodium*), which indicates the possibility that LECA relied on an overlapping set of genes in regulating its genome. Broadly, the method presented here – Pheno2Seq – can be applied to characterize gene expression in other microbial species that are marked by cryptic life cycle transitions as well as to identify gene expression patterns of cells fixed *in situ* as a means of asking about changing gene expression across ecological and temporal gradients.

## Materials and Methods

### Culture acquisition and maintenance

We used *Allogromia laticollaris* Cold Spring Harbor (CSH) for all experiments. Cultures were obtained and maintained as described in Timmons et al. (9). Cells were kept in ventilated tissue culture flasks with 32-ppt solution of Volvic water and instant ocean sea salt (Instant Ocean) in an incubator on a 12 hour dark/light cycle and fed *Isochrysis* algae (Carolina Biological). Algal cultures were maintained in ventilated culture flasks and fed on F/2 medium (UTEX, Austin, TX).

### Fixing with Glyoxal and staining with Hoechst

We pipetted cells from culture flasks into 200 µL of instant ocean water, then fixed a cell pellet (∼50 cells) in 200 µL of 3.2% glyoxal solution pH 6.0, and incubated at +4 ℃ for 20 minutes. The glyoxal solution is based on the protocol in Channathodiyil & Houseley (29), consisting of 124 µL of deionised water, 40 µL ethanol, 2 µL acetic acid (Thermo Fischer, A38S500) and 16 µL of 40% glyoxal (Sigma-Aldrich, St. Louis, MO 63103, US. 50649-25ML). We adjust the pH to 6 with sodium hydroxide. After 20 minutes, the fixative is rinsed off with 200 µL of 1X PBS (Sigma-Aldrich, Saint Louis, MO. P3813). Cells are permeabilized with 200 µL of 100% ice-cold methanol for 10 minutes on ice and rinsed in 1X PBS. We stain cells in 100 µL of 0.01 µg/ml Hoechst 33342 (Thermo Fisher Scientific, Waltham, MA. 62249) and incubate at room temperature on a shaker table for two hours. Hoechst is rinsed off with 1X PBS and we image cells within 24 hours. All PBS rinses are three washes and all steps are performed in a 1.5 mL centrifuge tube.

### Imaging

To collect images (light and fluorescent), we suspend stained cells in 1 mL of 1x PBS in glass bottomed Petri dishes (Fisher scientific, Hampton, NH) and capture images using a Leica TCS SP5 Laser Scanning Confocal Microscope. To prevent excessive laser damage (a concern for downstream RNA analysis), we captured a single cross-section image on 1024 x 1024 resolution with an acquisition speed of 200 Hz and line average of 8. The UV laser with excitation of 405 enables capture of light emitted from Hoechst between 425-562 nanometers (nm).

### Life stage identification

We characterized the life stage of each *Allogromia laticollaris* cell while imaging based on morphological criteria and nuclear stage (9). Cells were identified as juvenile (cell diameter <100um), adults (>100um) or schizonts (offspring visible within the parental test), and then characterized as uninucleate, multinucleate or *Zerfall*. The nuclear stage was determined by direct visualization of Hoechst stained nuclei under the confocal microscope. We photodocumented with confocal imaging (Data S1A) then picked single cells for transcriptome amplification. In total, we have 8 juveniles (3 uninucleate, 5 multinucleate), 11 adults (5 uninucleate, 6 multinucleate) and 3 *Zerfall* cells.

### Whole transcriptome amplification of image specimens

Immediately following image acquisition, we froze cells according to the protocol for cDNA library preparation with TaKaRa SMART-Seq (Cat 634940, 69). We purify the libraries with two rounds of AMPure XP beads and check libraries with fragment analysis. We ran PCRs with foraminifera-specific primers to validate that we captured *Allogromia* RNA. We prepped libraries for Illumina sequencing on the Novaseq platform with the Nextera XT DNA library preparation kit (Illumina). Once sequences were back, we checked read quality with FastQC (Babraham Bioinformatics, version 0.12.0). Illumina adapters were removed and filtered with a quality score of 24 with BBmap (70). Transcriptomes were *de-novo* assembled with rnaSpades (71).

### Assigning transcripts to gene families and building trees with EukPhylo

We used EukPhylo (version 1.0) to assign gene families (GFs) to assembled sequences (26, 72). Gene families were determined based on BLAST similarity to ∼15k gene families referred to as the “hook”. We further analyzed gene families that we identified by hand as having terms related to meiosis, genome dynamics, and chromatin organization according to annotation in OrthoMCL version 6 (Data S1D). Finally, we included a robust list of 485 highly expressed gene families across *Allogromia* in this study and 39 other foraminifera curated per Katz et al. (26) plus epigenetic gene families defined in Weiner et al. (73).

Based on presence in *A. laticollaris,* we used EukPhylo part 2 to build 1426 gene trees (Data S1D, fig. S2), with a Guidance (Version 2.02) cutoff of 0.3, allowing us to retain diverse taxa (74). We compiled a list of outgroup taxa for tree building, including 891 non-foram outgroup taxa, 39 non-*Allogromia* foraminifera, plus the 26 novel *Allogromia* transcriptomes in this study (Data S1A). Trees were built with IQTree (version 2.4.0, LG+Gamma substitution model and the fast mode), a lenient model given the stringency of Guidance alignment (75).

### Curation with phylogenomics

We used EukPhylo’s “clade-grabbing” tools to select sequences in trees as a means of building a set of reference *Allogromia* sequences for analyses of function and differential expression. To ensure we proceeded with *Allogromia* sequences rather than biological contaminants (e.g cocultures), we curated sequences extensively. First, we selected all *Allogromia* subclades from the gene trees that had seven foraminiferal sequences, at least two of which were the *Allogromia* sequenced for this study, resulting in 1689 subclades. We inspected any resulting small clades to remove sequences that we suspected represented contaminants based on their position within the original taxon-rich phylogenies (Data S1C). To remove sequences that only differed in their ends (where we have less confidence), we trimmed 30 base pairs from the ends of all sequences then clustered nucleotide sequences at 99% identity with cd-hit (76) (cd-hit -d 0 -c 0.99). Finally, we manually curated those with many recent paralogs to remove chimeras, ultimately resulting in 1881 sequences in our reference transcriptome (see next paragraph).

To identify putative chimeras in generating our final reference list, we used a Python script that processed each gene family separately. For each fasta file, we filtered sequences by length in three rounds to remove fragments and assembly artifacts. First, we removed sequences shorter than 67% of the median length of the alignment, plus we removed sequences bigger than 150% of the recalculated mean length. Finally, we retain only sequences within 75-125% recalculated mean length of the remaining sequences. Sequences of sufficient length were then classified as paralogs or alleles (recent duplicates) through an iterative procedure, with the aim of distinguishing legitimate sequences from chimeras. First, the sequence with highest “fit-score” (product of assembly coverage and sequence length, parsed from sequence headers) is designated as the master sequence and pairwise-aligned to all remaining sequences using MAFFT. For each alignment, we calculated sequence divergence in sliding windows (75bp, step size 1 bp), excluding windows with 10 or more gaps. We classified a pair as alleles (and removed from further iterations) if mean divergence lower than 4% and no window in the central 80% of the alignment exceeded 20% of divergence. Otherwise the sequence was retained as a paralog for the next iteration. We repeat rounds of master sequences and paralogs definition, until no new paralogs are identified or only one sequence remains. We then grouped paralogs into families based on shared sequence identity. For each pairwise alignment among retained paralogs, we identified the longest consecutive identical alignment, and sequences that shared at least 75 bp were assigned to the same family; we visualized these sequences in pairwise divergence plots and sliding windows analyses to look for, and remove, putative chimeras. For the histone paralogs, we also built a custom coulson plot to visualize the paralogs including expression information (Fig. 4) and we used MEGA 12 to estimate evolutionary divergence between retained histone paralogs (fig. S1) (77–79).

### Differential gene analysis

We aligned and quantified all raw reads from the 35 *Allogromia* to our reference sequences with RSEM and pseudoaligned and quantified the same data with Salmon. Both methods gave similar results, we proceeded with the RSEM quantification. We excluded all cells with <750,000 reads that were not *Zerfall* (n= 8) since we believe this to be part of the biology of the *Zerfall* stage. We performed quality control and exploratory analysis on the raw reads, to assess sequencing depth and expression variance across samples, using plotPCA and ggplot2 functions in R. We filtered lowly expressed genes prior to the gene expression analysis using a counts-per-million (CPM >= 1, in at least 2 samples) criteria implemented by edgeR package.

We estimated size factors using median-of-ratios normalization to library size normalization and correct for sequencing depth differences. We used the DESeq2 package in R (33, 34, 80) to perform differential expression analysis between biologically sequential life stages established in (9) (Fig. 1). We estimated surrogate variables using the SVA package based on the normalization counts to control for unwanted variations between the conditions (80). Gene-wise dispersion was estimated under a negative binomial generalized linear model, and differential expression was assessed using Wald tests (DeSeq2). We adjusted p-values for multiple testing using the Benjamini–Hochberg FDR method (adjusted p-values = padj). Genes were considered differentially expressed if they had an absolute log2fold change of at least 1 and an adjusted p-value of 0.05 or lower. We visualized differentially expressed genes using volcano plots and MA plots using EnhancedVolcano and pheatmap packages. Overlap between differentially expressed gene sets was assessed using Venn diagrams. All scripts and configuration files for the analysis are available in Github (https://github.com/Katzlab/Pheno2Seq).

### Functional analysis

We annotated protein-coding sequences derived from the reference transcripts using Blast2GO, combining BLAST searches using the UniProt/Swiss-Prot database with InterProScan protein family domain predictions. Each of these databases generate GO terms annotations (27, 81) that we merged to generate a gene-GO mapping that we used for the enrichment analysis. We conducted Gene Ontology enrichment in R, using both over-representation analysis (ORA) and gene set enrichment analysis (GSEA). We used clusterProfiler (82), enrichplot (83) and fgsea packages (84) for this enrichment analysis. We performed the analysis for all GO terms, and for Biological Process, Molecular Function and Cellular Component categories separately. We excluded gene sets smaller than 10 differentially expressed genes from this analysis to avoid bias. We visualized enriched terms using dot plots and categories–gene network plots (cnetplots). We visualized GSEA results using normalized enrichment score (NES) bar plots. To reduce redundancy, remove animal related annotations and facilitate our interpretation of the enrichments, we clustered the enriched terms by hierarchy on the GO structure. We tested different hierarchical levels on this clustering analysis. All scripts and configuration files for the analysis are available in Github (https://github.com/Katzlab/Pheno2Seq).

## Acknowledgments

We are grateful to Judith Wopereis (Smith College) for help with confocal microscopy, to Xyrus Mauer-Alcala for early discussions, to Tejasvi Kumaran for illustration in Figure 2, and to many Katzlab members for their support. Funding for this project includes two National Science Foundation grants to L.A.K. DEB 2439030 and DEB-2230391.

## Author Contributions

Conceptualization: LAK; Methodology: LAK, EGS, ML, GR; Investigation: EGS, GR, LAK, ML; Visualization: GR, EGS; Funding acquisition: LAK; Project administration: LAK; Supervision: LAK; Writing – original draft: EGS, GR, LAK; Writing – review & editing: EGS, GR, LAK, ML

## Supplementary Materials

Figs. S1 to S2

Data S1 to S10

