## Supplemental Figures 1 and 2 for "Elucidating changes in gene expression driving nuclear dynamics across a foraminiferal life cycle with Pheno2Seq"

**The PDF file includes:**

Supplementary Text
Figs. S1 to S2
References

**Other Supplementary Materials for this manuscript are on FigShare include the following:**

Figshare link:
https://figshare.com/projects/Pheno2Seq_analysis_of_differentially_expressed_genes_in_i_Allogromia_laticollaris_i_/268976
Data S1 to S10

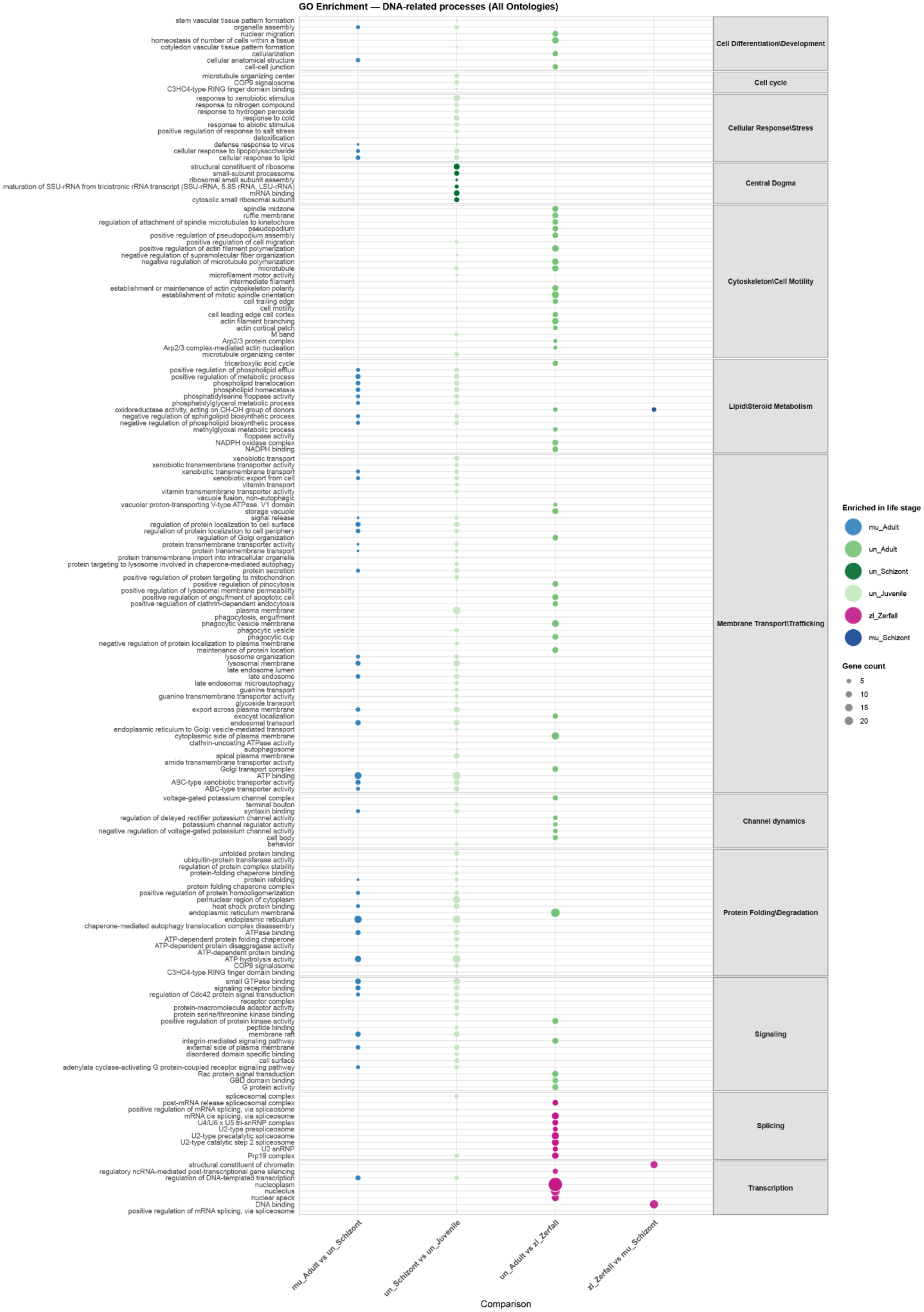
GO Enrichment — DNA-related processes (All Ontologies)
stem vascular tissue pattern formation
organelle assembly
nuclear migration
homeostasis of number of cells within a tissue
cotyledon vascular tissue pattern formation
cellularization
cellular anatomical structure
cell-cell junction
Cell Differentiation\Development
microtubule organizing center
COP9 signalosome
C3HC4-type RING finger domain binding
Cell cycle
response to xenobiotic stimulus
response to nitrogen compound
response to hydrogen peroxide
response to cold
response to abiotic stimulus
positive regulation of response to salt stress
detoxification
defense response to virus
cellular response to lipopolysaccharide
cellular response to lipid
Cellular Response\Stress
structural constituent of ribosome
small-subunit processome
ribosomal small subunit assembly
maturation of SSU-rRNA from tricistronic rRNA transcript (SSU-rRNA, 5.8S rRNA, LSU-rRNA)
mRNA binding
cytosolic small ribosomal subunit
Central Dogma
spindle midzone
ruffle membrane
regulation of attachment of spindle microtubules to kinetochore
pseudopodium
positive regulation of pseudopodium assembly
positive regulation of cell migration
positive regulation of actin filament polymerization
negative regulation of supramolecular fiber organization
negative regulation of microtubule polymerization
microtubule
microfilament motor activity
intermediate filament
establishment or maintenance of actin cytoskeleton polarity
establishment of mitotic spindle orientation
cell trailing edge
cell motility
cell leading edge cell cortex
actin filament branching
actin cortical patch
M band
Arp2/3 protein complex
Arp2/3 complex-mediated actin nucleation
microtubule organizing center
Cytoskeleton\Cell Motility
tricarboxylic acid cycle
positive regulation of phospholipid efflux
positive regulation of metabolic process
phospholipid translocation
phospholipid homeostasis
phosphatidylserine floppase activity
phosphatidylglycerol metabolic process
oxidoreductase activity, acting on CH-OH group of donors
negative regulation of sphingolipid biosynthetic process
negative regulation of phospholipid biosynthetic process
methylglyoxal metabolic process
floppase activity
NADPH oxidase complex
NADPH binding
Lipid\Steroid Metabolism
xenobiotic transport
xenobiotic transmembrane transporter activity
xenobiotic transmembrane transport
xenobiotic export from cell
vitamin transport
vitamin transmembrane transporter activity
vacuole fusion, non-autophagic
vacuolar proton-transporting V-type ATPase, V1 domain
storage vacuole
signal release
regulation of protein localization to cell surface
regulation of protein localization to cell periphery
regulation of Golgi organization
protein transmembrane transporter activity
protein transmembrane transport
protein transmembrane import into intracellular organelle
protein targeting to lysosome involved in chaperone-mediated autophagy
protein secretion
positive regulation of protein targeting to mitochondrion
positive regulation of pinocytosis
positive regulation of lysosomal membrane permeability
positive regulation of engulfment of apoptotic cell
positive regulation of clathrin-dependent endocytosis
plasma membrane
phagocytosis, engulfment
phagocytic vesicle membrane
phagocytic vesicle
phagocytic cup
negative regulation of protein localization to plasma membrane
maintenance of protein location
lysosome organization
lysosomal membrane
late endosome lumen
late endosome
late endosomal microautophagy
guanine transport
guanine transmembrane transporter activity
glycoside transport
export across plasma membrane
exocyst localization
endosomal transport
endoplasmic reticulum to Golgi vesicle-mediated transport
cytoplasmic side of plasma membrane
clathrin-uncoating ATPase activity
autophagosome
apical plasma membrane
amide transmembrane transporter activity
Golgi transport complex
ATP binding
ABC-type xenobiotic transporter activity
ABC-type transporter activity
Membrane Transport\Trafficking
voltage-gated potassium channel complex
terminal bouton
syntaxin binding
regulation of delayed rectifier potassium channel activity
potassium channel regulator activity
negative regulation of voltage-gated potassium channel activity
cell body
behavior
Channel dynamics
unfolded protein binding
ubiquitin-protein transferase activity
regulation of protein complex stability
protein-folding chaperone binding
protein refolding
protein folding chaperone complex
positive regulation of protein homooligomerization
perinuclear region of cytoplasm
heat shock protein binding
endoplasmic reticulum membrane
endoplasmic reticulum
chaperone-mediated autophagy translocation complex disassembly
ATPase binding
ATP-dependent protein folding chaperone
ATP-dependent protein disaggregase activity
ATP-dependent protein binding
ATP hydrolysis activity
COP9 signalosome
C3HC4-type RING finger domain binding
Protein Folding\Degradation
small GTPase binding
signaling receptor binding
regulation of Cdc42 protein signal transduction
receptor complex
protein-macromolecule adaptor activity
protein serine/threonine kinase binding
positive regulation of protein kinase activity
peptide binding
membrane raft
integrin-mediated signaling pathway
external side of plasma membrane
disordered domain specific binding
cell surface
adenylate cyclase-activating G protein-coupled receptor signaling pathway
Rac protein signal transduction
GBD domain binding
G protein activity
Signaling
spliceosomal complex
post-mRNA release spliceosomal complex
positive regulation of mRNA splicing, via spliceosome
mRNA cis splicing, via spliceosome
U4/U6 x U5 tri-snRNP complex
U2-type prespliceosome
U2-type precatalytic spliceosome
U2-type catalytic step 2 spliceosome
U2 snRNP
Prp19 complex
Splicing
structural constituent of chromatin
regulatory ncRNA-mediated post-transcriptional gene silencing
regulation of DNA-templated transcription
nucleoplasm
nucleolus
nuclear speck
DNA binding
positive regulation of mRNA splicing, via spliceosome
Transcription
mu_Adult vs un_Schizont
un_Schizont vs un_Juvenile
un_Adult vs zl_Zerfall
zl_Zerfall vs mu_Schizont
Comparison
Enriched in life stage
mu_Adult
un_Adult
un_Schizont
un_Juvenile
zl_Zerfall
mu_Schizont
Gene count
5
10
15
20

**Fig. S1.**

Bubble plot summarizing significantly enriched GO terms (Biological Process, Molecular Function, and Cellular Component combined) identified from pairwise differential expression comparisons between life cycle stages in *Allogromia*. Each row corresponds to an enriched GO term, and terms were grouped in broad functional categories (gray boxes). Columns represent the four pairwise comparisons with significant differential expression changes. Dot color indicates the life cycle stage in which the term is enriched, and dot size is proportional to the number of genes differentially expressed that contain the GO term, according to the lateral legends. Only GO terms meeting the significance threshold (p-adust < 0.05) in at least one comparison are shown. GO terms describing animal/plant specific structures or processes not applicable to our study organism (e.g., photosynthesis, synapse) were removed prior to plotting when possible.

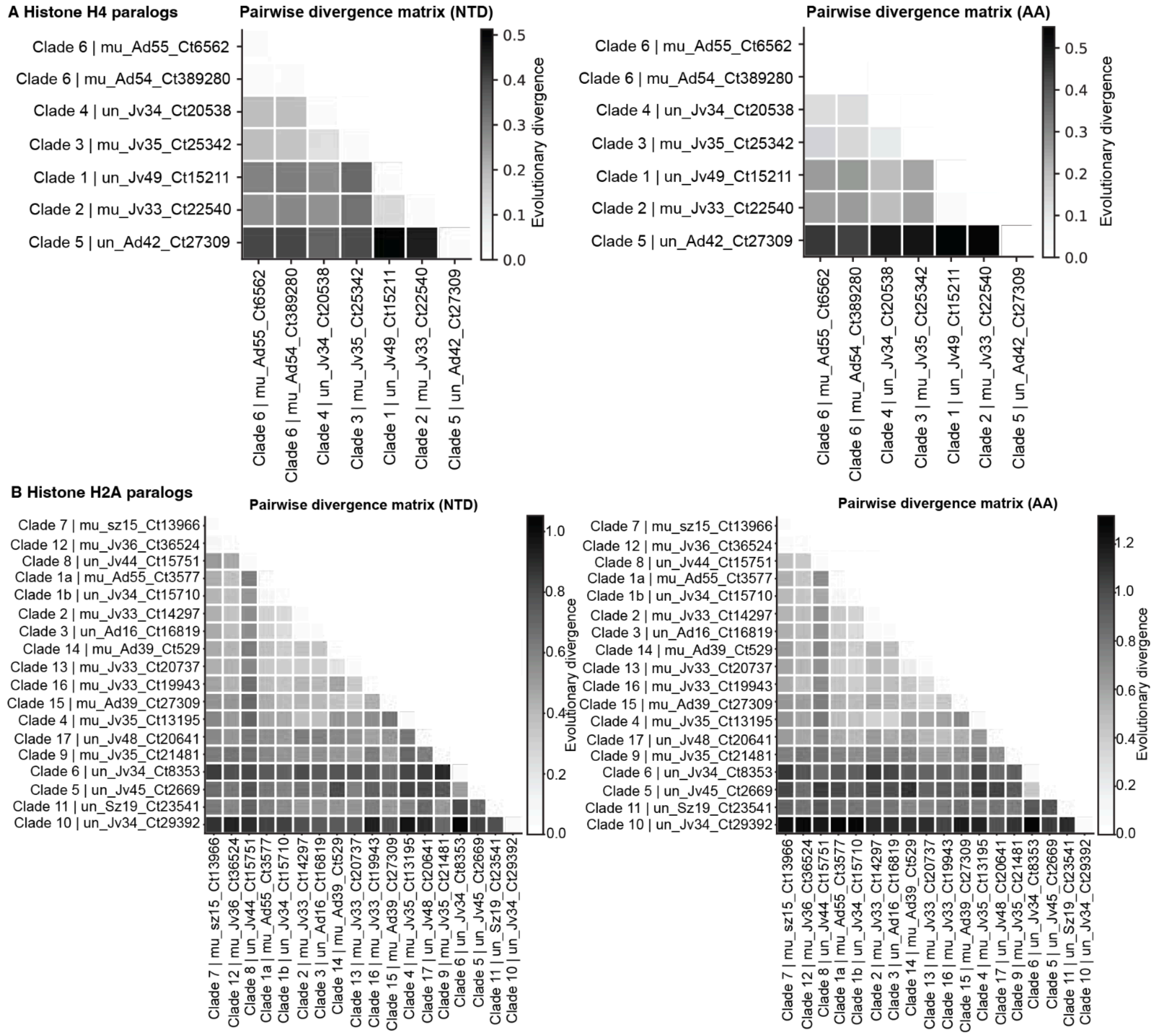


**Fig. S2.**

Pairwise evolutionary divergence among histone H4 and H2A paralogs. (A) Histone H4 paralogs. Pairwise divergence matrices for seven H4 paralog sequences retained in our study, labeled by clade (Fig.4) and sample of origin (e.g., mu_Ad55_Ct6562), calculated from nucleotide (NTD, left) and amino acid (AA, right) sequences. Rows and columns are ordered by similarity, darker cells indicate greater evolutionary divergence between the sequence pair (scale bar at right from each matrix). (B) Histone H2A paralogs. Pairwise divergence matrices for eighteen H2A paralog sequences retained in the study, labeled and ordered as in (A). Estimates of Evolutionary Divergence between Sequences. The number of amino acid and base substitutions per site between sequences is shown. For the nucleotide (NTD) analyses we used the Maximum Composite Likelihood model (*1*). The analytical procedure encompassed coding nucleotide sequences using 1st, 2nd, 3rd, and non-coding positions. The pairwise deletion option was applied to all ambiguous positions for each sequence pair resulting in final sets comprising (A, left) 342 positions for Histone H4 alignments and (B, left) 762 positions for Histone H2A alignments. For the amino acid (AA) analyses we used the Poisson correction model (*2*). The analytical procedure encompassed amino acid sequences. The pairwise deletion option was applied to all ambiguous positions for each sequence pair resulting in a final data set comprising (A, right) 114 positions for Histone H4 alignments and (B, right) 254 positions for Histone H2A alignments. Evolutionary analyses were conducted in MEGA12 (*2*, *3*).

**Data S1A. (All Cells in study)**

Cells in the study with life cycle stage metadata and sequencing summary statistics. Summary of samples used in this study, including number of reads sequenced, life cycle stage (e.g., schizont, adult), nuclear type (uni = uninucleate, multi = multinucleate), cell diameter (µm), quantified (whether the sample passed quality filtering and was included in downstream quantitative analyses), image label (identifier linking the sample to its corresponding microscopy image), number of sequences (NSeqR2G) and number of OGs (NOGsR2G) recovered after EukPhylo processing, GC content at third codon position variation width (WidthGC3) and cell type used for grouping/plotting (e.g., "schizont_multi", "adult_uni").

**Data S1B. (All taxa in study)**

Taxon sampling and taxonomy for gene trees in this study. List of species included to generate the gene trees in this study using the EukPhylo pipeline. Code is the internal identifier used by the pipeline, encoding higher level taxonomic grouping and species (e.g., Pl_gr_Atha for *Arabidopsis thaliana* within the "Pl" [Plantae]/"gr" green plants group); Taxonomy, full taxonomic lineage from supergroup to genus, as retrieved from NCBI Taxonomy; Species, full binomial species name (e.g., *Arabidopsis thaliana*).

**Data S1C. (clades removed)**

Manual curation summary of gene family subtrees used to identify contaminant or chimeric sequences before downstream gene expression analysis. If outgroup taxa were not as expected, the clade was removed from study. Subtree name (name of clade file output by Eukphylo clade grabbing); OG (OrthoMCL6 code); Clade (ID assigned by EukPhylo); Remove (clades manually marked for removal).

**Data S1D. (final OG list in reference)**

List of OrthoMCL orthologous genes (OG6) retained in the final dataset after paralogs curation, manual curation and quality filtering. Each row corresponds to one ortholog group used in downstream expression and phylogenetic analyses.

**Data S2. (All image captured for study as .TIFF)**

Raw microscopy images collected for this study, provided as individual .TIFF files.

**Data S3. (.fasta of all sequences in study)**

File containing all sequences in the study, provided as .fasta files.

**Data S4. (All parental trees in study)**

File containing all parental trees in the study, provided as .tre files.

**Data S5. (Paralog/alleles_result_files)**

File containing all the results for our paralog/allele analysis used to curate our reference.

**Data S6. (.fasta of all reference sequences in study)**

File containing all retained reference sequences in the study, provided as .fasta files.

**Data S7. (DE_genes_ALL)**

Differential gene expression results across pairwise life cycle stage comparisons. Differential expression (DE) results from pairwise comparisons between life cycle stages and nuclear conditions, computed using DESeq2. Genes were considered significantly differentially expressed at a 1 fold change and an adjusted p-value (padj) < 0.05. (A) Summary tab. Comparison (name of the pairwise comparison); Description (life cycle stage transitions represented); GroupA (base group) /GroupB (test group), groups compared; Total_genes (number of genes tested); DE_genes (number of significantly differentially expressed genes); Upregulated_A/Downregulated_A (number of DE genes with higher/lower expression in Group A relative to Group B). (B-I) Per-comparison tabs ([Comparison]_DE_genes_ALL), SeqName (gene identifier); GroupA/GroupB (groups compared); baseMeanA/baseMeanB/baseMean (mean normalized expression in Group A, Group B, and overall); log2FoldChange (log2 fold change of Group A relative to Group B); lfcSE (standard error of the log2 fold change); stat (Wald test statistic); pvalue (raw p-value); padj (Benjamini-Hochberg adjusted p-value); Description (direction of differential expression).

**Data S8A. (Sterner_readmapping_data_pres_absence)**

Gene family expression and differential expression summary across life cycle stages. For each reference sequence (RefSeqName) we assigned an OrthoMCL orthologous gene group (OG6, Gene family). Reference sequences are listed with its classification as a differentially expressed gene (DEG) or low-abundance transcript (Rare) (Type_gene). The Conditions_regulated column lists the pairwise comparisons in which the transcript was significantly differentially expressed and the direction of regulation (UP/DOWN) in each case. The seven subsequent columns color by the life cycle stage in which we found the reference sequence upregulated as color palette in Fig1. We provided the functional annotation based on OrthoMCL ortholog group assignment (OrthoMCL annotation) and BLAST similarity to the NCBI non-redundant protein database (NCBI annotation); for OrthoMCL annotations we provided multiple alternatives considering that many times the result was ambiguous (Alternative 2–5 OrthoMCL annotation). Then we provided the median expression across all samples (Median expression), and individuals replicate raw expression values, labeled by cell type and assigned number in our internal database.

**Data S8B. (Sterner_readmapping_data_pres_absence)**

Histone paralogs expression and differential expression summary across life cycle stages. Column descriptions are as in Data S8A.

**Data S8C. (Eggnog mapper output)**

Sequences for reference transcripts were analyzed by Eggnog mapper (*4*) to infer gene properties (e.g., gene name, PFams, EC numbers)

**Data S9. (gene_category_assignments_final_EDIT)**

Dominant functional category assignment for each reference sequence based on GO term annotation. For each reference sequence (RefSeqName), we report its expression classification (Type_gene: Constitutive, DEG, or Rare) as defined in Data S7A. For building the plots in Fig.1 and giving a general view of the functional categories assigned, we counted the total number of GO terms annotated to the sequence (N_GO_terms), and the number of distinct broader functional categories (N_Categories) represented among those GO terms. The single functional category with the highest representation among a gene's annotated GO terms was assigned as its Dominant_Functional_Category; genes with no clear majority category, or whose top categories were tied, are flagged in Is_Tie (TRUE/FALSE). Other_Avoided indicates whether the "Other" category was excluded when resolving ties or assigning dominance (TRUE/FALSE), so that a more informative category was assigned preferentially over "Other" where possible.

**Data S10A. (Supplementary_Enriched_GOterms)**

Gene ontology (GO) enrichment analysis of the differentially expressed genes across the pairwise life cycle stages comparisons. For each pairwise comparison (Group A/Group B; e.g., un_Adult vs. zl_Zerfall), differentially expressed genes in the indicated direction (Regulation: UP or DOWN), were tested for enrichment of GO terms (ID, Description) within the three GO term categories (Comparison_Type), Biological Process (BP), Molecular Function (MF), or Cellular Component (CC), and the three combined (ALL). The DNA_related column flags GO terms associated with DNA-related processes (used in Fig. 3), and Category assigns each term to a broader functional grouping (e.g., Nuclear Organization/Chromatin, Protein Folding/Degradation, Transcription) used for cluster results. The enrichment analysis report the following statistics: the number of differentially expressed genes annotated with the GO term relative to the total number of differentially expressed genes tested in that comparison (GeneRatio), and the corresponding ratio for the background gene set (BgRatio); The proportion of background genes annotated with the term that were recovered among the differentially expressed genes (RichFactor); The ratio of the observed to expected proportion of genes annotated with the term (FoldEnrichment); The deviation of the observed gene count from that expected by chance (zScore). The raw p-value (pvalue) and after multiple-testing correction using both the Benjamini-Hochberg method (p.adjust) and the q-value approach (qvalue) statistical significance. The identifiers (RefSeqName) of all differentially expressed genes contributing to enrichment of the term (geneID), and the total number of such genes (Count).

**Data S10B. (DNA related Supplementary_Enriched_GOterms)**

Gene ontology (GO) enrichment analysis of the differentially expressed genes across the pairwise life cycle stages comparisons flagged as DNA_related in S10A. Column descriptions are as in Data S10A.
